# A two-compartment system for subfreezing temperatures preservation of large volumes of organic matter in an isochoric system

**DOI:** 10.1101/2022.08.31.506083

**Authors:** Gabriel Năstase, Florin Botea, George-Andrei Beșchea, Ștefan-Ioan Câmpean, Alexandru Barcu, Irinel Popescu, Boris Rubinsky, Alexandru Șerban

## Abstract

This is a technology paper on the design of and test results from an 11-liter isochoric (constant volume) chamber, for the preservation of large, organs in a supercooled state. Details of the design are given, as well as a proof that the chamber is isochoric. Five repeats show that in this large chamber, ice nucleation of steam distilled water occurs at – 2 °C within less than 12 hours, in all the repeats. An examination of the experimental results suggests that the ice nucleation starts on the inner walls of the isochoric chamber. A new two compartment isochoric chamber was designed to reduces the probability of ice nucleation on the walls of the chamber. In the two-compartment system, the biological matter and the preservation fluid are introduced in a sealed low-density polyethylene bag, and placed in the center of the isochoric chamber, in such a way that the bag does not touch the walls. The space between the inner walls of the isochoric chamber and the outer walls of the bag are filled with a fluid with a composition that does not freeze at the storage temperature. Three repeat experiments with steam distilled water and with *in vitro* pig liver show that with this technique, the system remained supercooled, without any ice nucleation for the duration of the experiments. Experiments were voluntary terminated at 48 hours of supercooling. This new technology may hold promise for long term preservation of large biological organs in a supercooled state, without the use of any chemical additives.

## INTRODUCTION

Preservation of biological matter in a supercooled thermodynamic state is a way to reduce the metabolism to subfreezing temperatures without the detrimental formation of ice. There is growing interest in preservation of biological matter in a supercooled state for various application in life sciences and food sciences (1). The idea of preserving biological matter in a supercooled state is not new (2). A variety of techniques for supercooling preservation without ice were developed throughout the years. To list a few: emulsification (2), elevated pressure (3), electromagnetic fields (4), cryoprotectant solutions at temperatures at which they do not freeze (5) (6), antifreeze proteins (7), partial freezing to mimic survival of freeze tolerant species (8) which recently reported major achievements (9), deep-supercooling of large volumes with surface sealing with immiscible fluids (10) which also reported major achievements (11).

Isochoric (constant volume) supercooling is a relatively recent area of research in the field of preservation of biological matter in a supercooled state. The basic concept of isochoric preservation draws from fundamental thermodynamics (12). In a simplistic explanation, it is related to the fact that the density of ice Ih (hexagonal ice crystal) is lower than the density of water. Therefore, when ice begins to form in an isochoric system, the pressure in the interior of the chamber will increase (13). Thermodynamic analysis has shown that, at the eutectic about 50% of the volume remains unfrozen, and biological matter can be preserved in the unfrozen volume, without freezing (13). While isochoric freezing has also useful applications in food technology and medicine e.g. (14), (15), this paper deals with isochoric supercooling. Thermodynamics (16), and experiments (17),(18),(19) have shown that isochoric conditions stabilize the metastable supercooling. Preservation of biological matter by supercooling was also achieved (20). However, the experiments on isochoric supercooling were done with relatively small chambers, in the 100 ml range.

A first goal of this technological study was to design and test a large-scale isochoric supercooling chamber, suitable for preservation of large organs or large quantities of food products. The information in this paper should facilitate making of such an isochoric chamber by researchers interested in the field. Performing preliminary studies with the isochoric chamber we observed that the solution in this large chamber becomes unstable faster than anticipated from the earlier experiments with small chambers. Therefore, another goal of the study was to find ways to improve the stability of isochorically supercooled solutions in large isochoric chambers. A way to improve the stability was found through the use of a two compartments system. The two-compartment isochoric chamber is also described and tested.

## 2. MATERIALS AND METHODS

### 2.1. ISOCHORIC REACTOR

The isochoric reactor is built around an isochoric chamber with an internal volume of 11 liters. The chamber is designed to withstand a pressure of 8 bar. The reactor is instrumented with a pressure transducer and two temperature sensors positioned at two different heights inside the chamber, a solution filling port and an overflow port. Figure 1 shows the top and side images of the assembled isochoric reactor.

**Figure 1.**
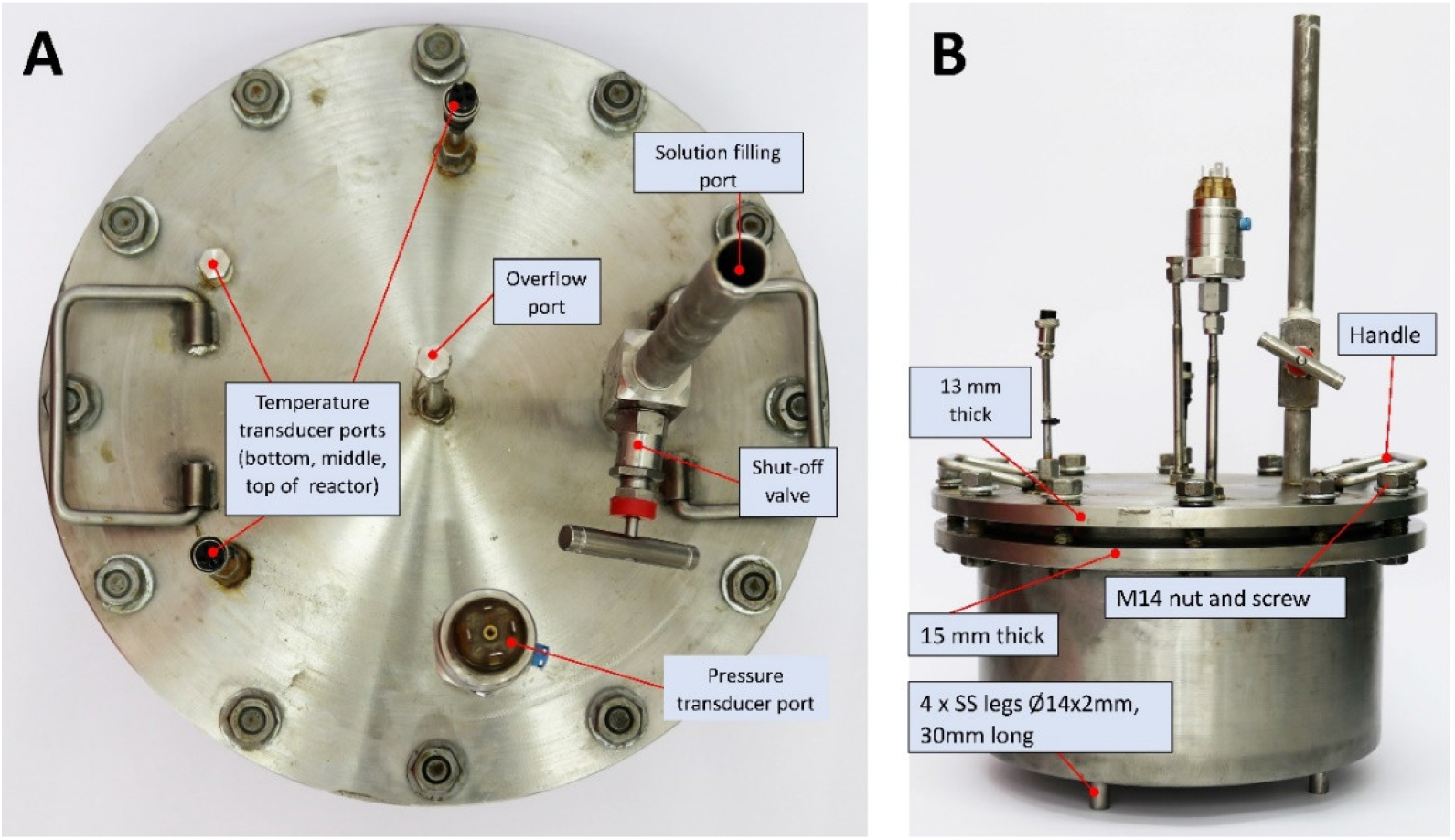
Top (A) and side (B) images of the assembled isochoric reactor

The heart of the isochoric reactor is the isochoric chamber. Figure 2 shows the top (A) and side view (B) of the open isochoric chamber. The view of the bottom of the isochoric chamber lid is in Figure 2C.

**Figure 2.**
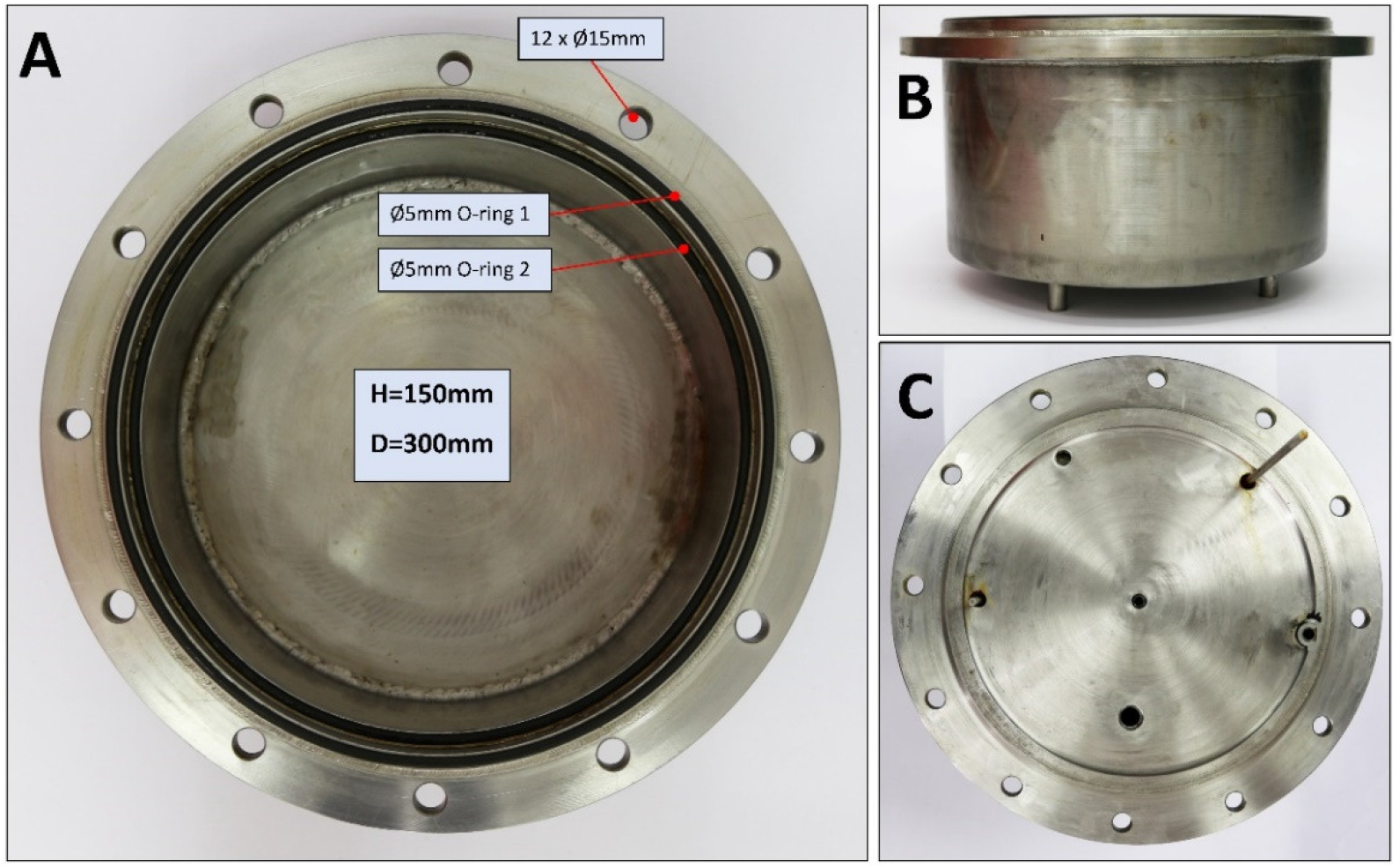
Top (A) and side view (B) of the open isochoric chamber. The view from the bottom of the isochoric chamber lid (C)

The isochoric chamber has an internal volume of 11 liters, and is designed to accommodate large organs. The main component of the chamber is a section of a DN300 pipe (austenitic stainless steel AISI 321 W1.4541 from ITALINOX ROMANIA SRL) with an internal diameter of 303 mm and a height of 150 mm. A 13 mm thick bottom plate of (austenitic stainless steel AISI 321 W1.4541 from ITALINOX ROMANIA SRL) was welded to the pipe. On the top of the tubular isochoric chamber is a 4×2mm indentation that can accommodate a rubber O-ring (Silicone rubber NBR70 O-ring, Dichtomatik - Freudenberg FST GmbH, Germany). A 10 mm height, 8 mm wide ring was welded to the exterior of the chamber at a distance of 10 mm from the top. The ring accommodated a second O-ring (Ø5mm Silicone rubber NBR70, designed for operating temperatures from -30°C to +100°C). Twelve holes in the ring accommodate the 16×90 screws. The reactor chamber lid is made of a 19 mm stainless steel (austenitic stainless steel AISI 321 W1.4541 from ITALINOX ROMANIA SRL) plate. The reactor cover is attached to the body of the reactor with twelve 16×90 screws, as shown in Figures 1 and 2. The reactor sits on four (Ø14×2mm) 30 mm height support legs (austenitic stainless steel AISI 321 W1.4541 from ITALINOX ROMANIA SRL) welded as shown in Figure 1. The purpose of these legs is to provide the necessary space to completely surround the isochoric reactor with the coolant (a mixture of water and ethylene glycol).

The isochoric reactor is instrumented with two temperature transducers, and a pressure transducer. Resistance temperature detectors RTD Endress-Hauser TR10 (Endress+Hauser AG, Switzerland) are used to measure the temperature at two different sites in the isochoric chamber, top and bottom. They can measure temperatures in the range from -200 °C to +600 °C in systems under pressures up to 75 bar (1088 psi). The pressure transducer is a Cerabar PMC11 gauge pressure transducer (Endress+Hauser AG, Switzerland) (Figure 3B). It features a capacitive, oil-free ceramic sensor and is able to measure gauge pressure from 400mbar up to 40bar. It should be noted that the system is designed for isochoric preservation by supercooling. A concern with isochoric supercooling chambers is that when the interior of the chamber accidentally begins to freeze, the interior pressure during isochoric freezing can increase rapidly, and destroy the chamber. The chamber described in this paper was designed to withstand a pressure of 8 bar. Obviously if the design pressure is chosen to be lower, it should have been possible to use lighter materials. As a safety measure the system can be equipped with a safety head and rupture disk or a safety valve for overpressure protection. Another safety measure involving temperature control will be discussed later.

**Figure 3.**
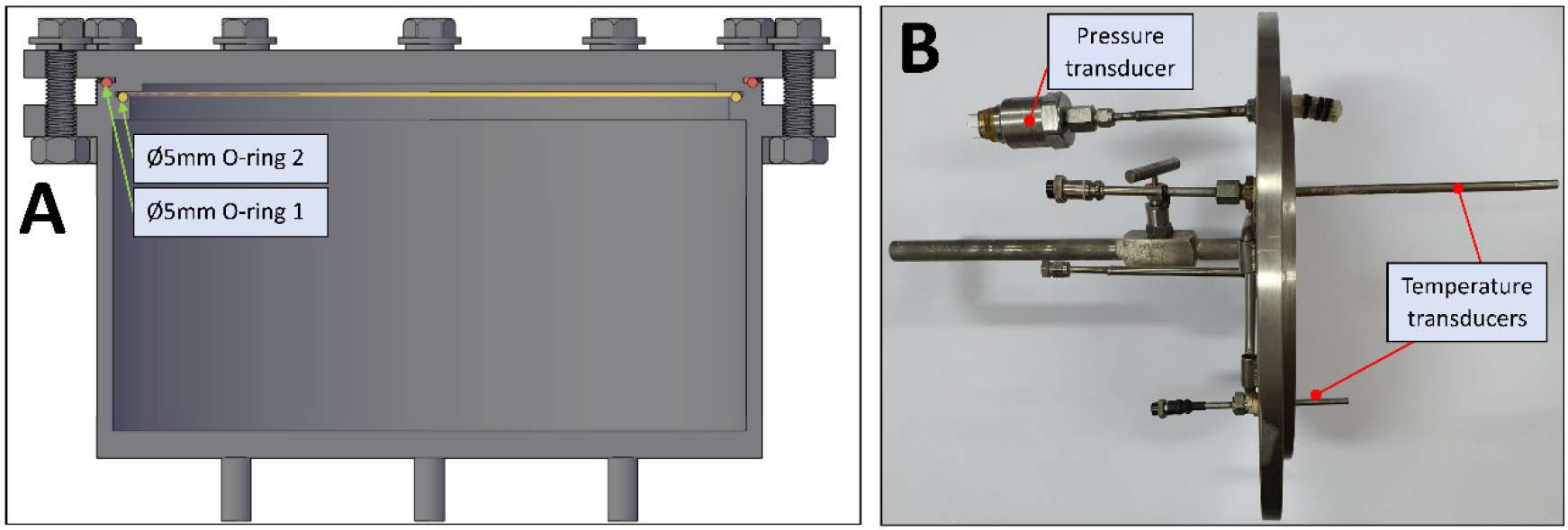
Vertical cross section through the margin of the vessel (A) and the assembled top plate, side view (B)

Figure 3A shows the assembly top plate and cylinder. The side view 3B shows the thermocouple connections and the pressure transducer. In addition, there is an overflow, air purging valve (Figure 1A). This is a very important part of the system as the isochoric concept fails when the system contains free air. The top surface of the cover plate also has a filling valve (Figure 1A). This is a BADOTHERM instrument needle valve (Badotherm BDTV910, maximum working pressure 413bar@38°C, 1/2”NPTm-1/2”NPTf, AISI316L, EU origin). This valve has a non-rotatable conical tip to ensure perfect alignment and all parts are made of AISI 316(L) low carbon stainless steel. Process connection (F) 1/2”NPT-m, instrument connection (F1) 1/2”NPT, maximum pressure 413 bar (6.000 psi) at 38°C and maximum temperature 240 °C. Figure 3B shows the thermistors on the interior part of the chamber. The two thermistors have different immersion lengths, 150 mm, and 50 mm.

Because all structures that protrude from to top surface can act as fins, they are insulated with thermal pipe insulation.

### 2.2. REFIGERATION SYSTEM

Refrigeration is an important part of the large volume isochoric supercooling system. The system requires a means to control the temperature of the entire chamber and to minimize fluctuations in temperature, inside the chamber. To this end we have designed a dedicated portable refrigeration system that can operate continuously for hours to maintain a volume of 11 liters in the isochoric chamber in a steady supercooled state. The schematic of the refrigeration system is shown in Figure 4.

**Figure 4.**
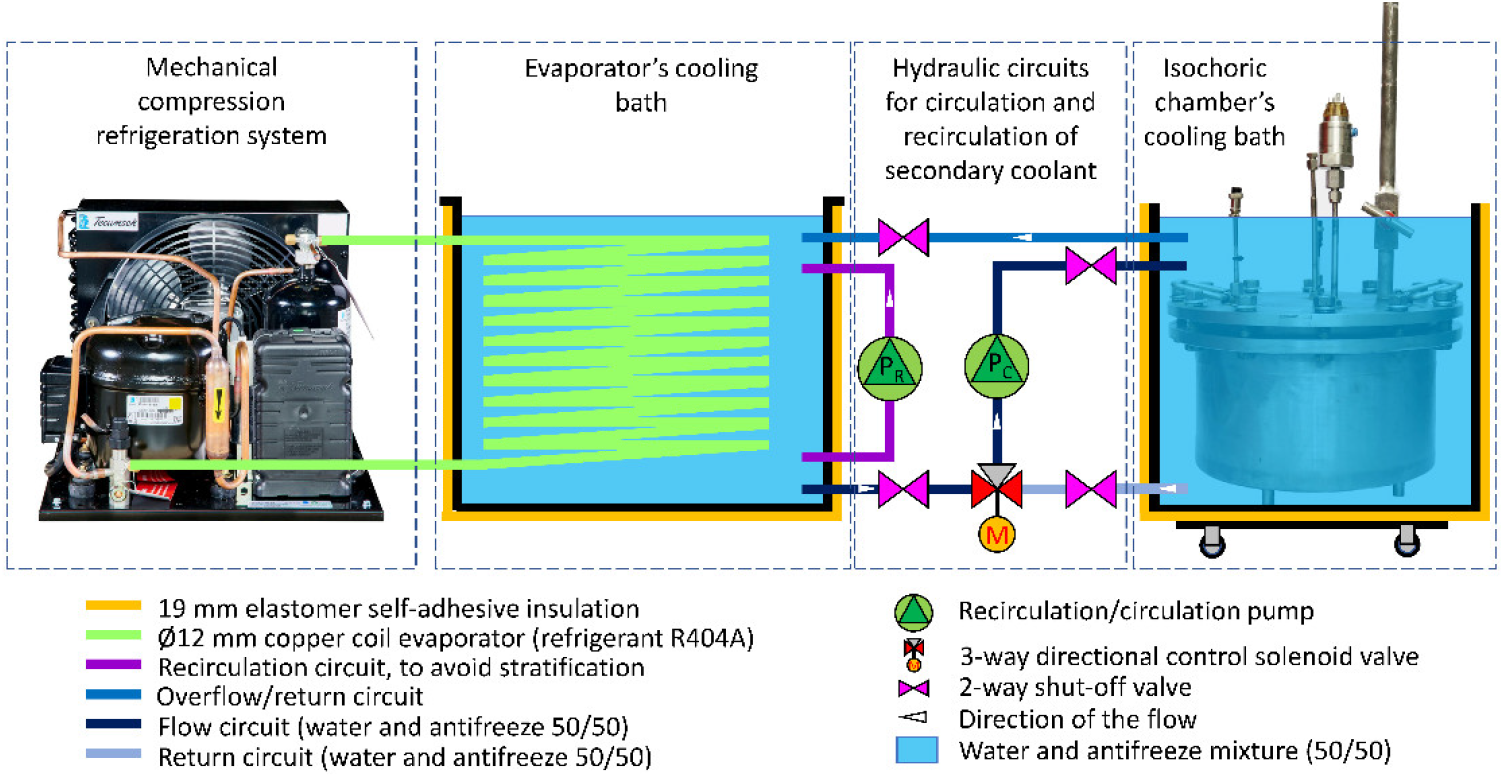
The schematic of the refrigeration system for large organs preservations

The refrigeration system consists of four elements. They are described from right to left. the first is an isochoric chamber cooling bath in which the isochoric chamber is completely immersed. The cooling bath is insulated and is connected to the second element. The second element is a hydraulic circuit for circulation and recirculation of a secondary coolant. This circuit is controlled with input from a temperature transducer in the reactor cooling bath and is designed to maintain a constant temperature in the cooling bath. The control system is also connected to the pressure transducer which stops the cooling the instant pressure elevation, (i.e. freezing) is recorded. The first two elements are required elements for the isochoric reactor. The second two elements, from right to left, are a typical recirculating cooling system which can be commercially purchased. We choose to build a dedicated refrigeration system. It consists of an evaporator cooling bath which supplies the coolant to the isochoric reactor cooling bath and the mechanical compression refrigeration unit. The assembled refrigeration system is shown in Figure 5.

**Figure 5.**
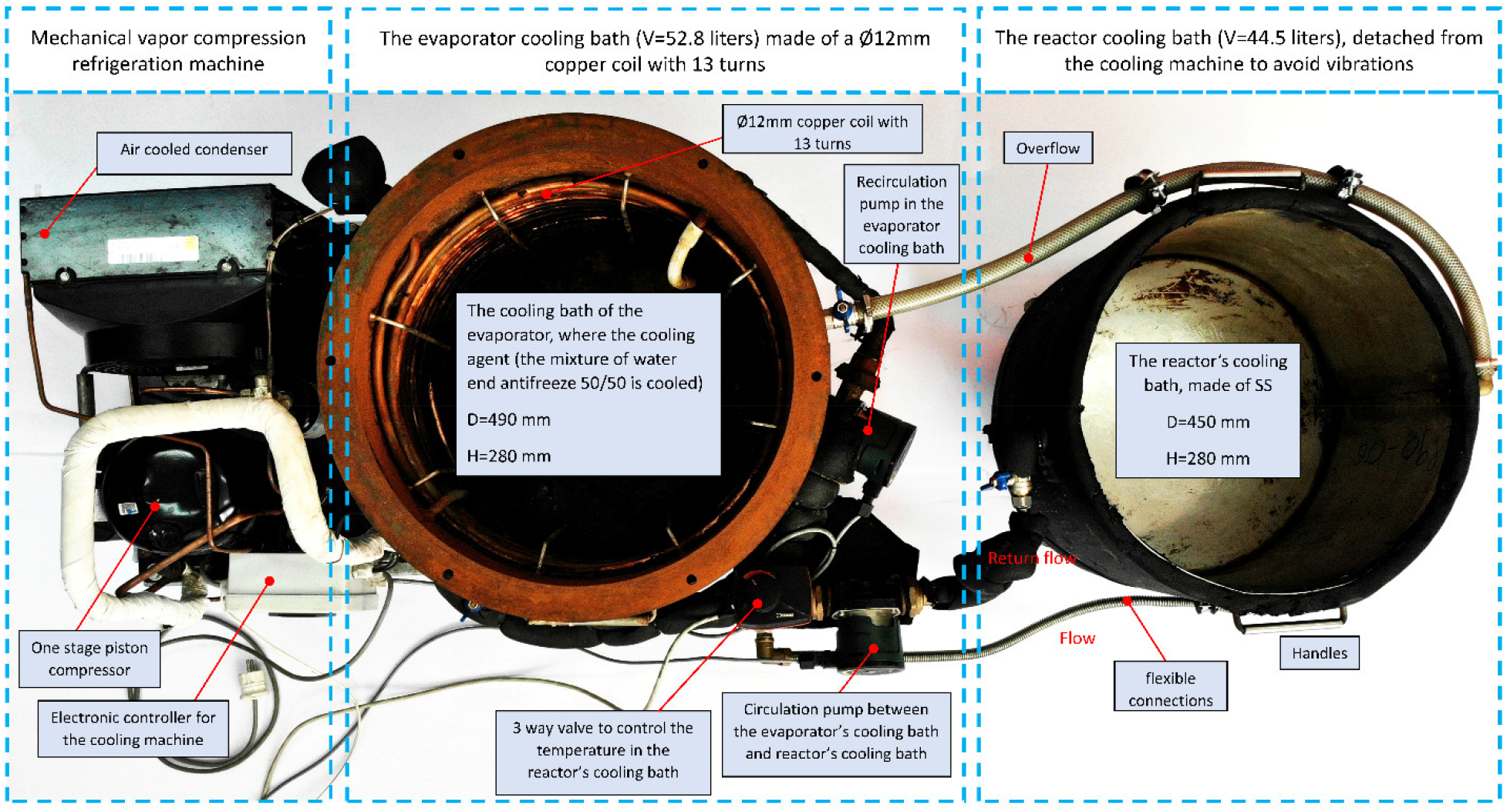
The assembled refrigeration system

Two refrigerator fluids are used. To cool the isochoric reactor, the bath system is using about 40 liters of 50/50 percent water and ethylene glycol. This fluid flows between the evaporator chamber and the cooling isochoric reactor chamber. The heart of the refrigeration system is a mechanical vapor compression cooling aggregate with an air-cooled condenser (Tecumsech AE4440 HR, PS30bar, TS125°C, working with a hermetic piston compressor Tecumsech AE-8036-BR). R404A is used as a refrigerant. To avoid stratification in the cooling bath we use a recirculation pump (DAB EVOSTA2, 40-70/130 1”, 3.6 m^3^/h with a maximum H of 7mH2O). A control system (Dixell, XR20XC, Emerson Climate Technologies GmbH, Germany) for the temperature inside the isochoric chamber receives inputs from the temperature and pressure sensors attached to the isochoric chamber. The control system turns on and off the flow through the isochoric chamber refrigeration circuit to maintain the desired temperature in the isochoric chamber. Figure 6A shows the isochoric reactor in the cooling chamber and Figure 6B shows the appearance of the reactor, immersed in the cooling chamber.

**Figure 6.**
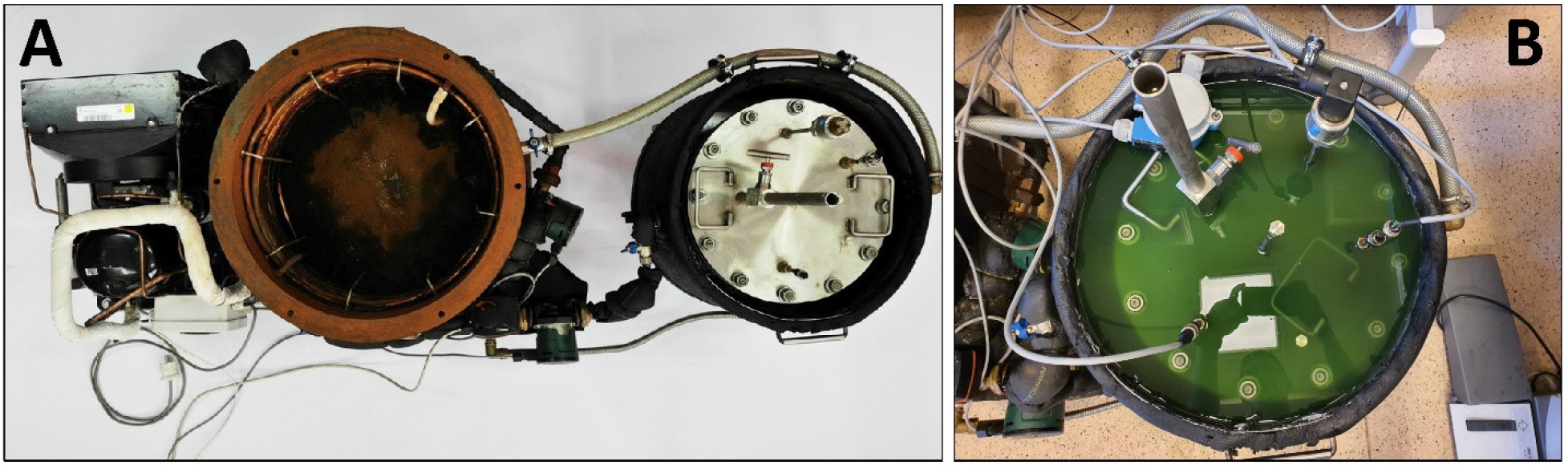
The isochoric reactor in the cooling chamber (A) and the appearance of the reactor, immersed in the cooling chamber (B)

### 2.3. CONTROL HARDWARE AND SOFTWARE

Figure 7A shows the entire ensemble with the control and automation unit. All devices and transducers are controlled from the control and automation panel in Figure 7A and Figure 7B.

**Figure 7.**
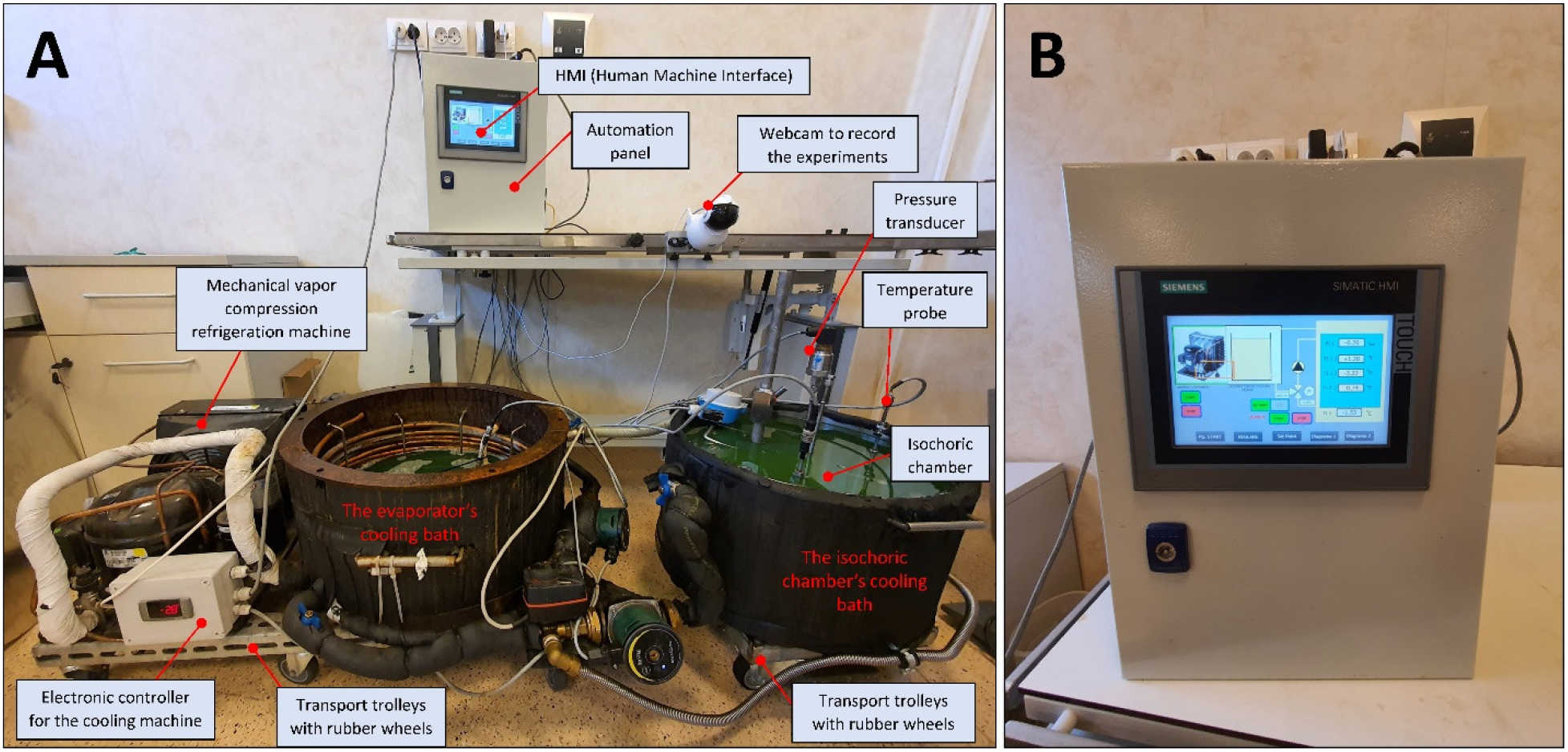
The entire ensemble with the control and automation unit (A) and the control and automation unit and the human-machine interface which has a touchscreen display (B)

The system has three parts: the programmable logic controller (PLC); sources, fuses, relays; and the Human-Machine Interface (HMI). The control panel allow users to control the process from distance via internet, using VNC and has the capability to store all information on memory stick or SD card. Figure 7B shows the control and automation unit and the human-machine interface which has a touchscreen display (Figure 7B). Particular control measures are the pressure transduced, the three temperature sensors in the isochoric reactor, the temperature transducer in the isochoric reactor bath. The control system controls the flow rate and the temperature in the isochoric reaction bath.

Figure 8 shows the various monitoring and control elements in the control and automation unit. Figure 9 shows the particular schematic of the connections for controlling the temperature of the fluid in the evaporator chamber. The insert shows a typical human interface display. Figure 10 shows the interactive screen display during the control of the evaporative chamber temperature (10A) and the isochoric reactor bath temperature (10 B).

**Figure 8.**
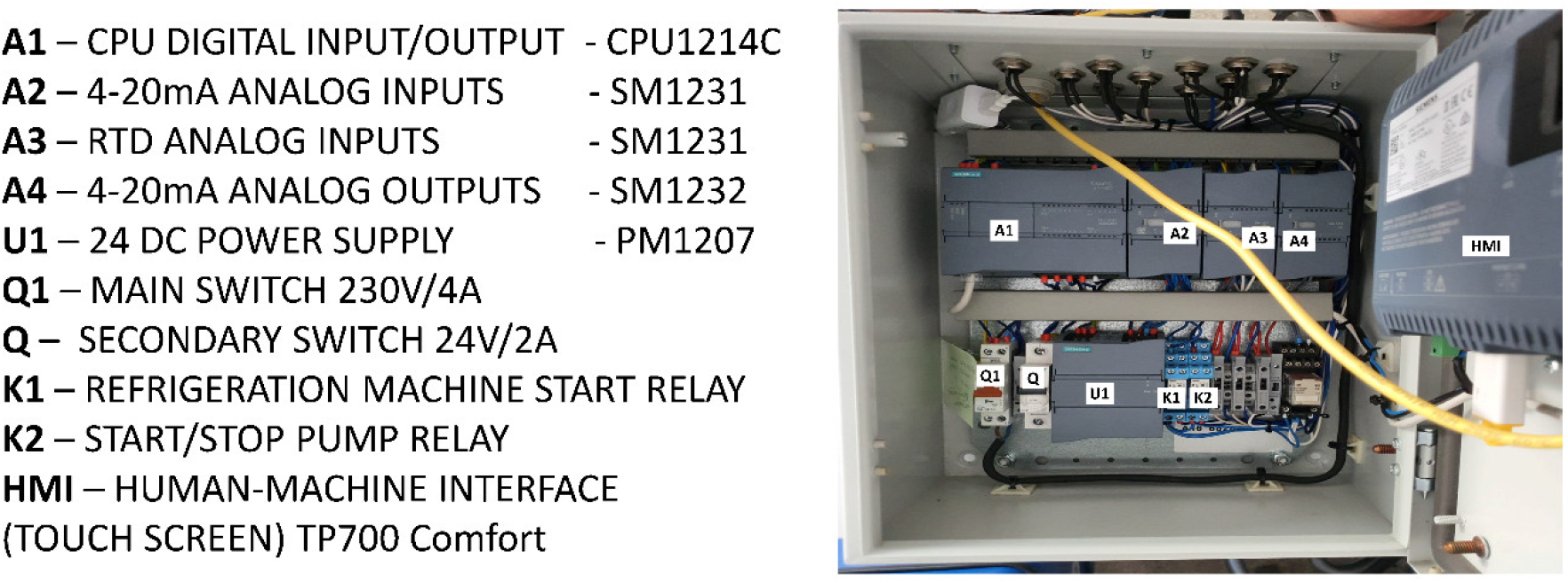
The various monitoring and control elements in the control and automation unit

**Figure 9.**
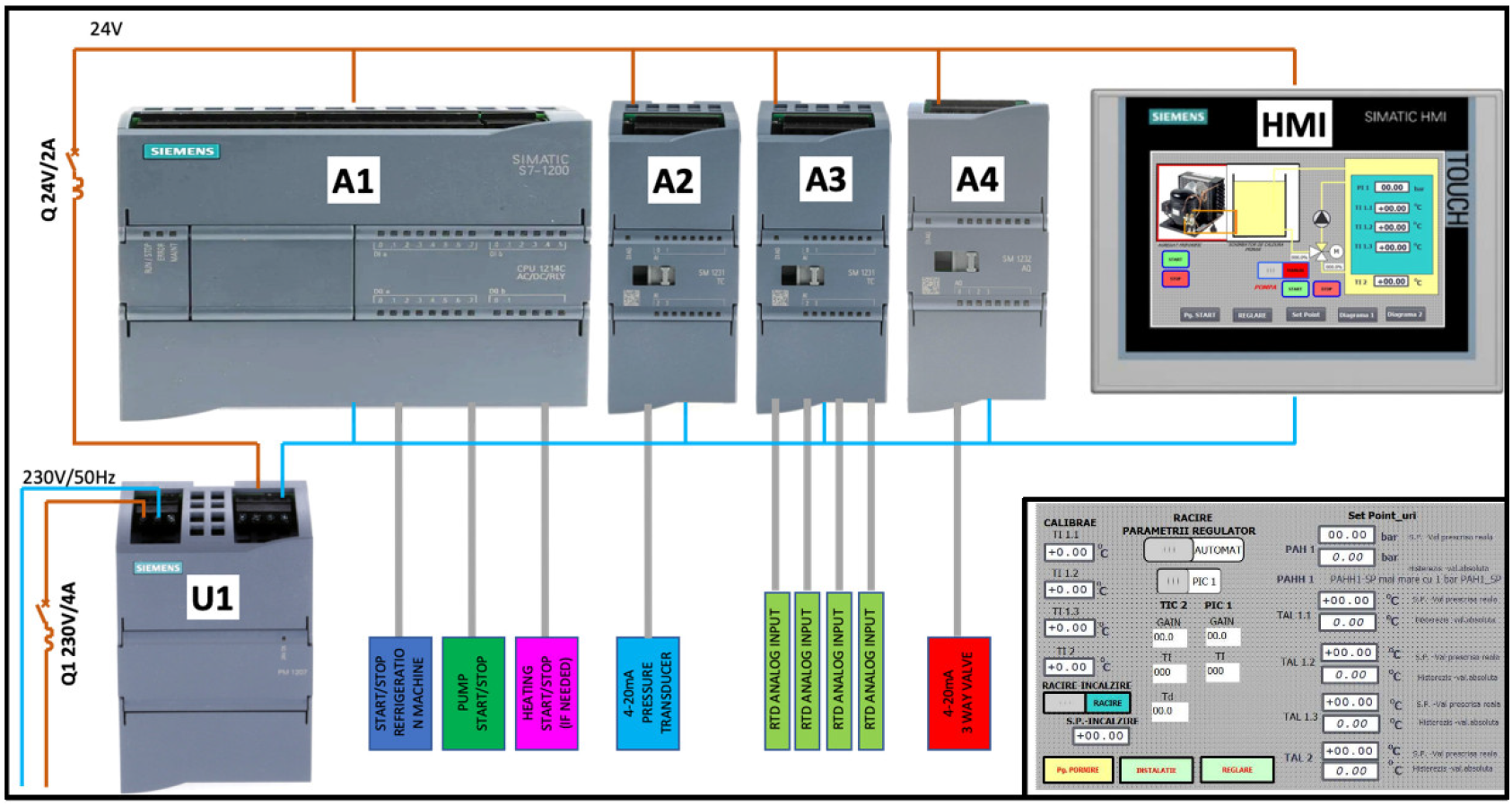
The particular schematic of the connections for controlling the temperature of the fluid in the evaporator chamber and the insert shows a typical human interface display

**Figure 10.**
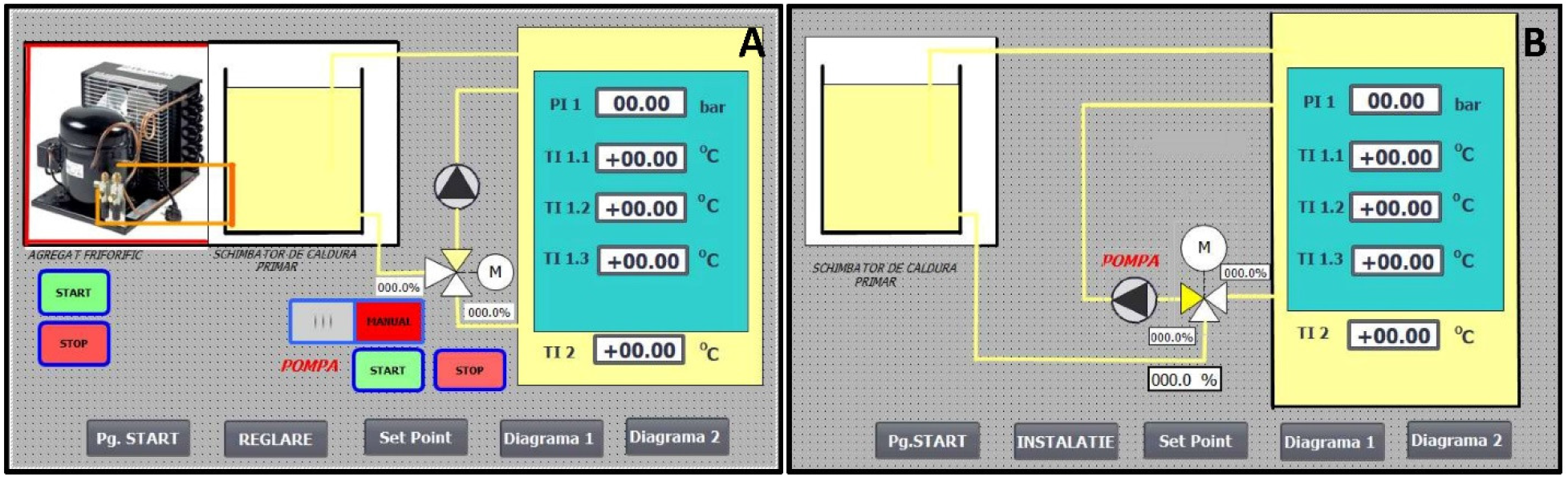
The interactive screen display during the control of the evaporative chamber temperature (A) and the isochoric reactor bath temperature (B).

The principle of operation of the system is as follows. The automation panel consists of a CPU (Siemens SIMATIC S7-1200 model CPU1214C) that can handle 14 digital inputs 24 V DC, 10 digital outputs 24 V DC/0.5 A and 2 analogue inputs 0-10 V. Three additional modules are mounted on the right side of the CPU to extend the digital or analog I / O capacity. The first two are of Siemens SM1231, one to take the signal from the pressure transducer and the second Siemens SM1231 is to take the signals from the RTDs. The last additional signal module Siemens SM1232 uses an output to control the three-way valve, that keeps the temperature constant in the reactor’s cooling bath. For power supply all components use a 24V switched mode power supply unit from Siemens (Siemens, PM1207). The human machine interface is provided using a Siemens Simatic HMI TP700 Comfort panel, with touch operation, 7” widescreen-TFT-display, 16 mil. colors, PROFINET interface, MPI/PROFIBUS DP interface, 12 mb user memory. This control panel allow users to control the process from distance via internet, using VNC (RealVNC® Limited, UK) and has the capability to store all information on memory stick or SD card. The human-machine interface has a touchscreen display where users can observe and interact with all the measured parameters in the system. The automation box includes analog inputs: 3 temperature gauges in the capsule with PT100 resistance temperature detectors - RTDs, 1 antifreeze temperature sensor, 1 absolute pressure gauge in the capsule, digital outputs: 1 refrigeration system command on relay K1, 1 antifreeze recirculation pump command in the reactor’s cooling bath on relay K2, analog outputs: 1 output for commanding the 3-way regulating valve. To record all experiments, we used an IP wireless smart camera (PNI, model IP930W 1080P) full HD 1080p with IR and night vision. The wireless connection to the system was made through a wireless router 4G LTE Cat6 (Huawei, model B525s-23a).

### 2.4. EXPERIMENTAL PROTOCOL

This section describes the assembly protocol and points to important steps in setting up the experiment. Errors in these steps may cause the failure of the experiments because of departure from isochoric conditions.

The first step is filling the isochoric reactor cooling bath and the evaporator chamber with a solution that can sustain the desired preservation temperature. In this study we have used a solution of 50% volume ethylene glycol, 50% volume water. In typical experiments this solution is precooled to the desired initial temperature. Because the thermal mass is large, precooling the system to the desired initial temperature can have an impact on the biological matter preservation protocol outcome.

The isochoric chamber is prepared as following. At the start of the experiment the correct placement of the O-rings and the secure placement of all the sensors are verified. After cleaning the interior of the isochoric chamber and the isochoric chamber top cover, to avoid any residuals on the walls and transducers, the open isochoric chamber is filled with the solution used in the experiment or application. Air bubbles may appear in the solution. It was shown earlier that the volume of free air in the system should be minimized and preferentially eliminated [(21)]. It is recommended to fill the isochoric chamber slowly and to overflow the chamber.

The next step is to place the lid carefully on the O-rings while the overflow valve on the top lid is kept open. The lid is secured to the body of the chamber using twelve screws and nuts as shown in Figure 3A.

As mentioned earlier free air in the isochoric system must be minimized and preferable eliminated. To this end, the overflow connection is opened after the lid was secured to the chamber. Then, fluid is added through the solution flowing port (Figure 1A) until it flows out from the overflow port. The valves to the two ports are closed and the screws are further tightened. At this stage it is important to monitor the pressure transducer, because the final tightening process can be used to preset the initial pressure in the experiments. This step is also crucial to the isochoric experiment because the initial pressure affects the thermodynamics of the process. There are two elements that must be carefully controlled when setting isochoric studies, the elimination of free air from the system and the initial pressure in the system. The next step is to immerse the isochoric reactor in the cooling bath.

The isochoric reactor temperature and pressure are controlled throughout the experiment and at the end of the prescribed period of storage the isochoric system is warmed to a temperature above freezing and opened.

## 3. RESULTS AND DISCUSSION

### 3.1 ICE NUCLEATION IN ISOCHORIC SUPERCOOLED SYSTEMS

Experiments were performed with the experimental setup and methods described in the material and methods section following the protocol described in the experimental protocol section. First, a study was performed with steam distilled water. The goal of this part of the study was to evaluate the stability of the supercooled fluid in a large isochoric chamber of about 11 liters. As mentioned earlier, previous experiments with isochoric supercooling were done with small volumes of fluid, of about 100 ml to 250 ml. To the best of our knowledge, this is the largest volume in which the stability of an isochoric supercooling fluid was ever tested. The test involved continuously measuring the pressure and temperature in a supercooled state. In all the experiments described in this paper, the average supercooling temperature was – 2°C.

According to our previous studies, albeit with much lower volumes, the supercooling of isochoric systems at – 2ºC should be stable, for days (18). It is important to note that in the studies reported in (18) the walls of the chamber were coated with a hydrophilic substance, petroleum jelly (Vaseline). In this study we have not coated the walls with Vaseline, because of technical difficulties with the uniform coating of such a large surface. In contrast to small volume experiments, in virtually all the experimental repeats with the large isochoric chamber (five), there was ice nucleation. It should be emphasized that in this study the fluid in the large isochoric chamber is exposed to continuous mechanical excitations due to the operation of the refrigeration motor and to temperature fluctuations induced by the on/off operation of the refrigeration system. Figure 11 shows the pressure in the isochoric chamber as a function of time. The measurements were taken at the discrete times listed in the figure. The figure shows that the experiment began at 14h27m11s. The pressure remained constant until 21h36m33s, for a period of 7h9m22s, after which it suddenly increased. This suggests that ice nucleation has occurred and also serves as proof that the system is isochoric. When ice forms in an isochoric system the pressure increases because the density of ice is higher than that of water. In closed isochoric systems, freezing leads to an increase in pressure (13), (22).

**Figure 11.**
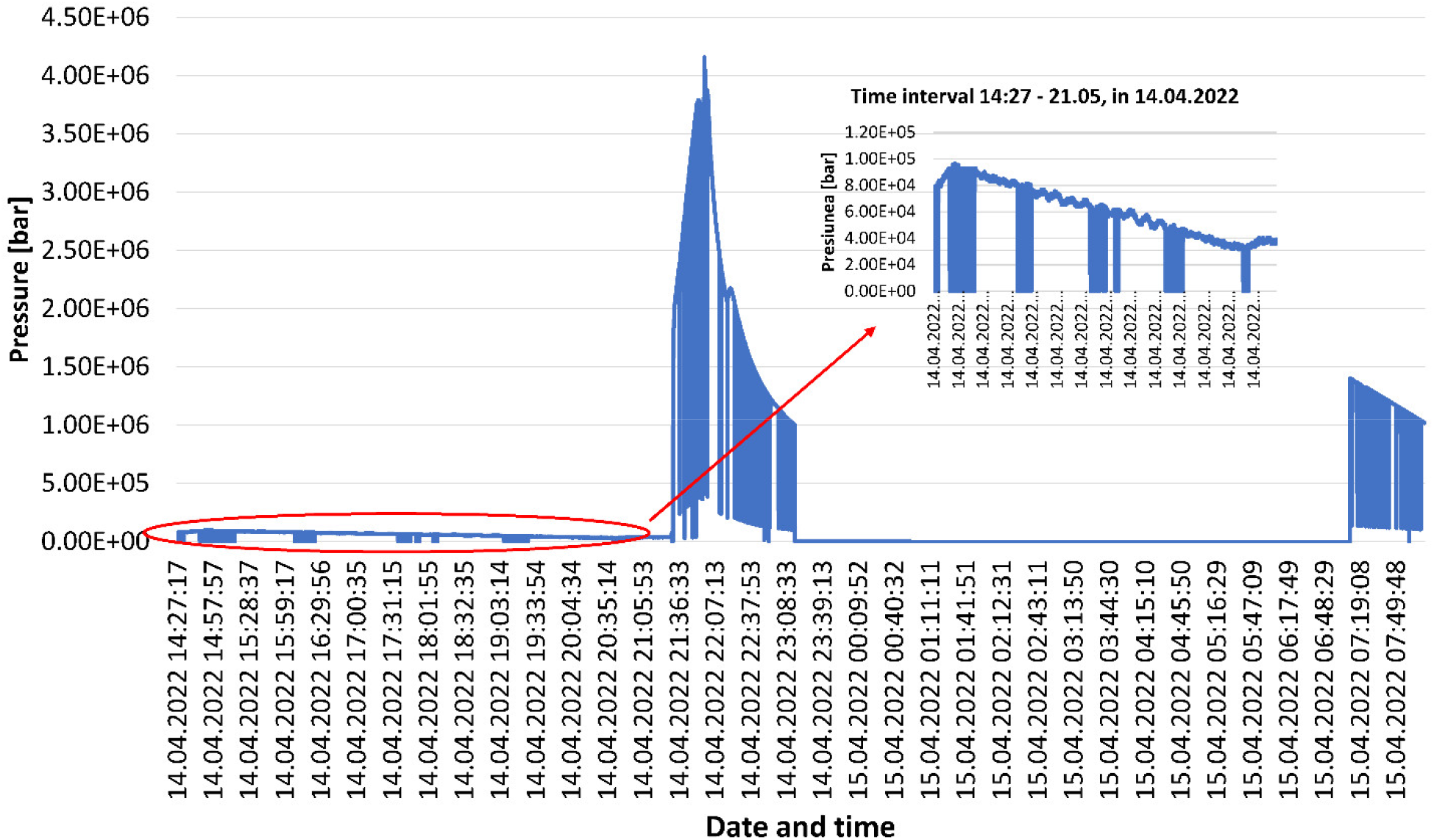
Pressure in the isochoric chamber as a function of time

Figure 12 shows the temperature at three sites inside the isochoric chamber top, middle and bottom) and in the fluid in the isochoric reactor cooling bath. The temperature recording began at 14h27m17s. The results show 21h36.33s the temperature measured by the upper thermistor suddenly began to increase. The temperature trace is typical to ice nucleation in a supercooled system. At the onset of nucleation, the temperature dropped slightly and then raised rapidly to the phase transition temperature.

**Figure 12.**
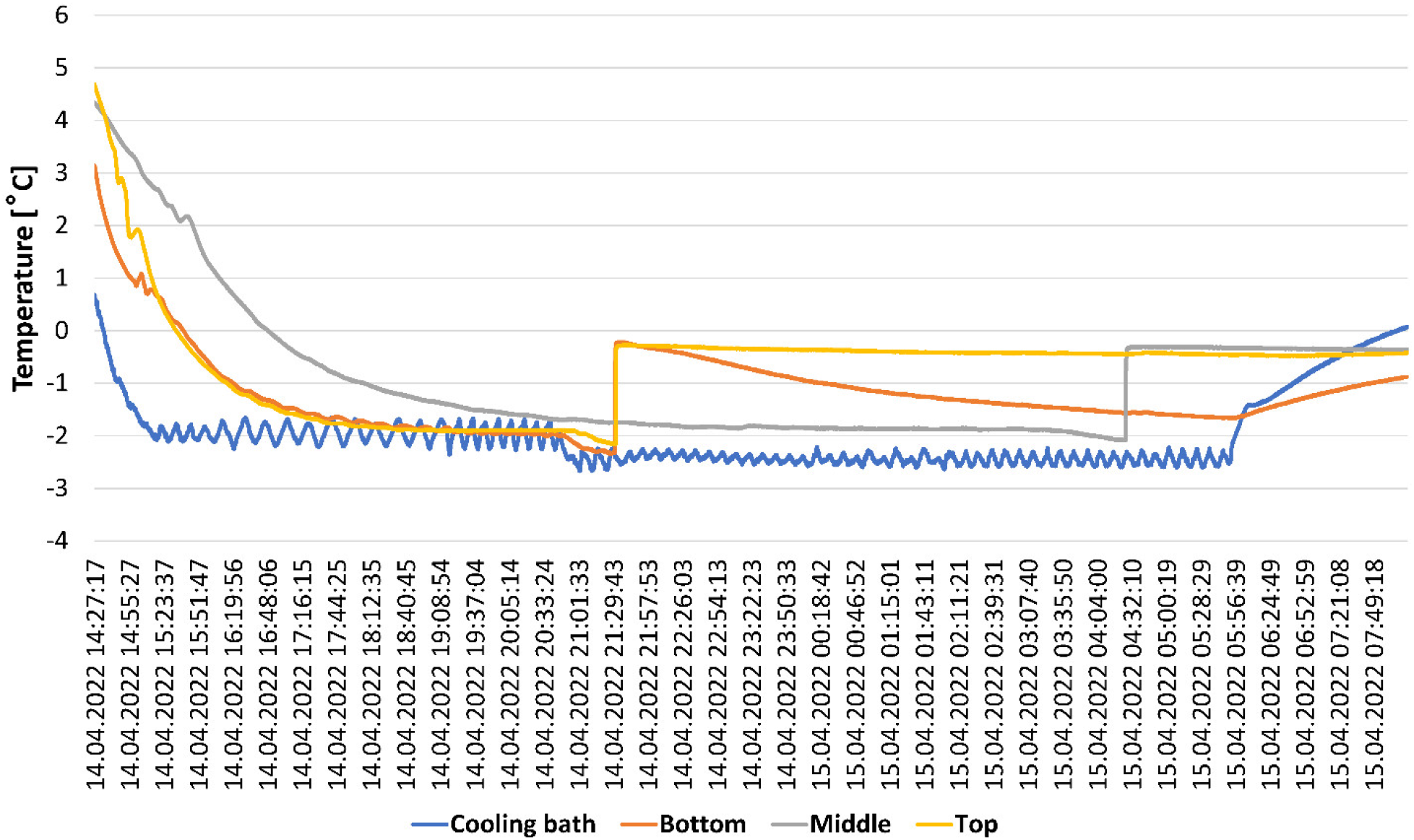
Temperature at three sites inside the isochoric chamber top middle and bottom) and in the fluid in the isochoric reactor cooling bath

The results in Figures 11 and 12 show that the sudden increase in pressure is associated with a simultaneous sudden increase in temperature. This is expected in an isochoric system. When ice begins to form in an isochoric system, the expansion of water as ice yields an increase in the pressure. The results of this part of the study can be viewed as proof that the experimental system is system is isochoric.

In summary, ice nucleation in the large isochoric supercooled chamber occurred much faster than anticipated (17) (18). The pattern of ice formation is shown in Figure 13. It is seen that the ice forms along the walls, in particular along the top surface and the regions with a weld. This tentatively suggests that the nucleation was induced by the isochoric chamber inner surface, which in this experiment is much larger than in previous experiments with smaller chambers and was not coated with a hydrophobic substance.

**Figure 13.**
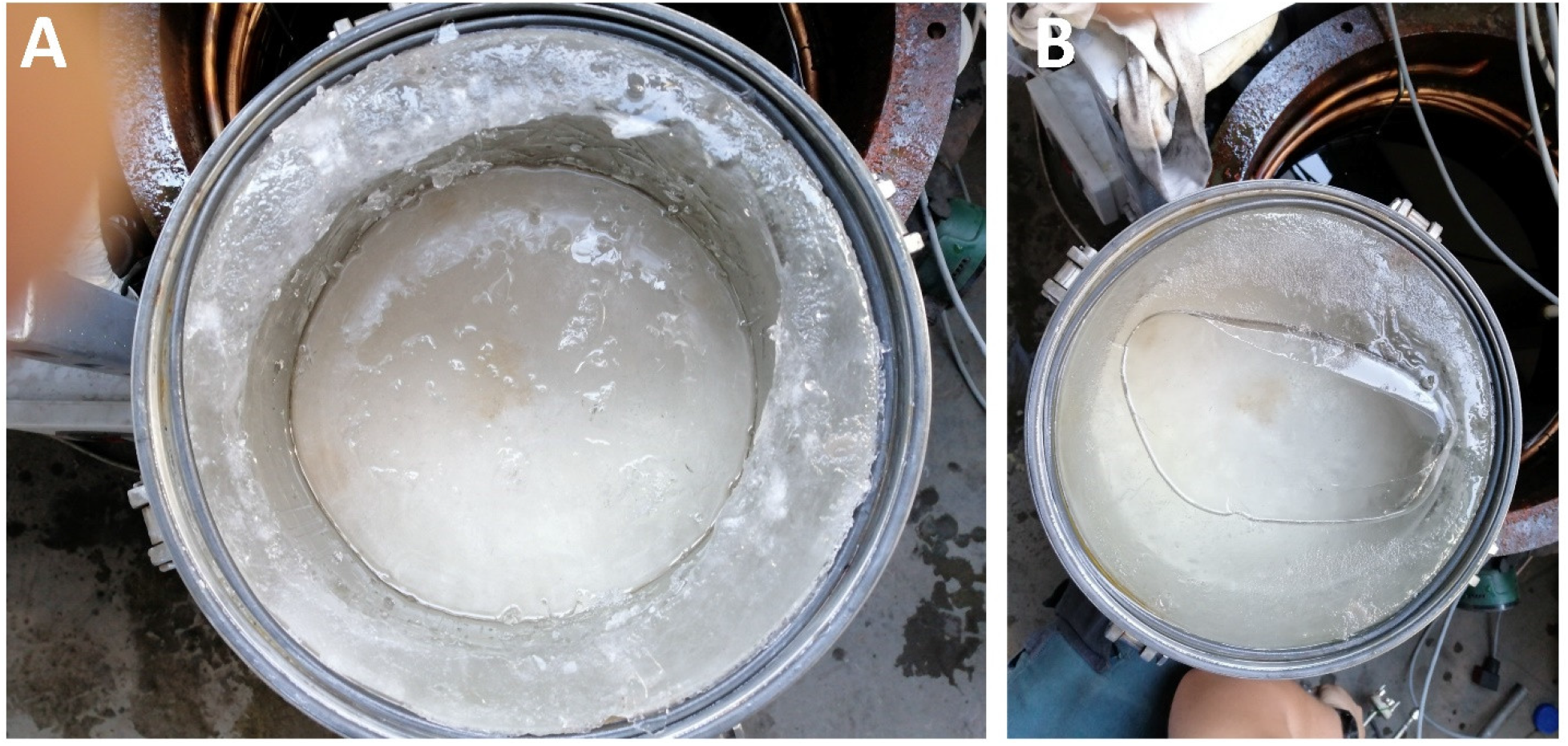
The pattern of ice formation

### 3.2 ELIMINATION OF ICE NUCLEATION ON THE INNER WALLS OF AN ISOCHORIC CHAMBER

A two-compartment design was developed and tested for long term isochoric supercooling by eliminating the possibility for ice nucleation along the inner walls of the isochoric chamber. To avoid nucleation from the isochoric chamber walls a sealable thin wall plastic bag made of light density polyethylene (LDP) is used as a second compartment. The LDP bag is filled with the fluid to be maintained in a supercooled state. For organ preservation, the organ with the preserving fluid is introduced in the bag. The bag is completely filled with the fluid with attention to avoid any air in the plastic bag. During a two-compartment isochoric supercooling preservation protocol, the plastic bag with the material to be preserved in a supercooled state is inserted in the isochoric chamber. The isochoric chamber is than filled, with a fluid with a freezing point lower than the isochoric supercooling storage temperature. Placing such a fluid between the inner walls of the chamber and the outer walls of the plastic bag will eliminate the possibility that nucleation will occur on the inner walls of the isochoric chamber. Figure 14 illustrates the concept.

**Figure 14.**
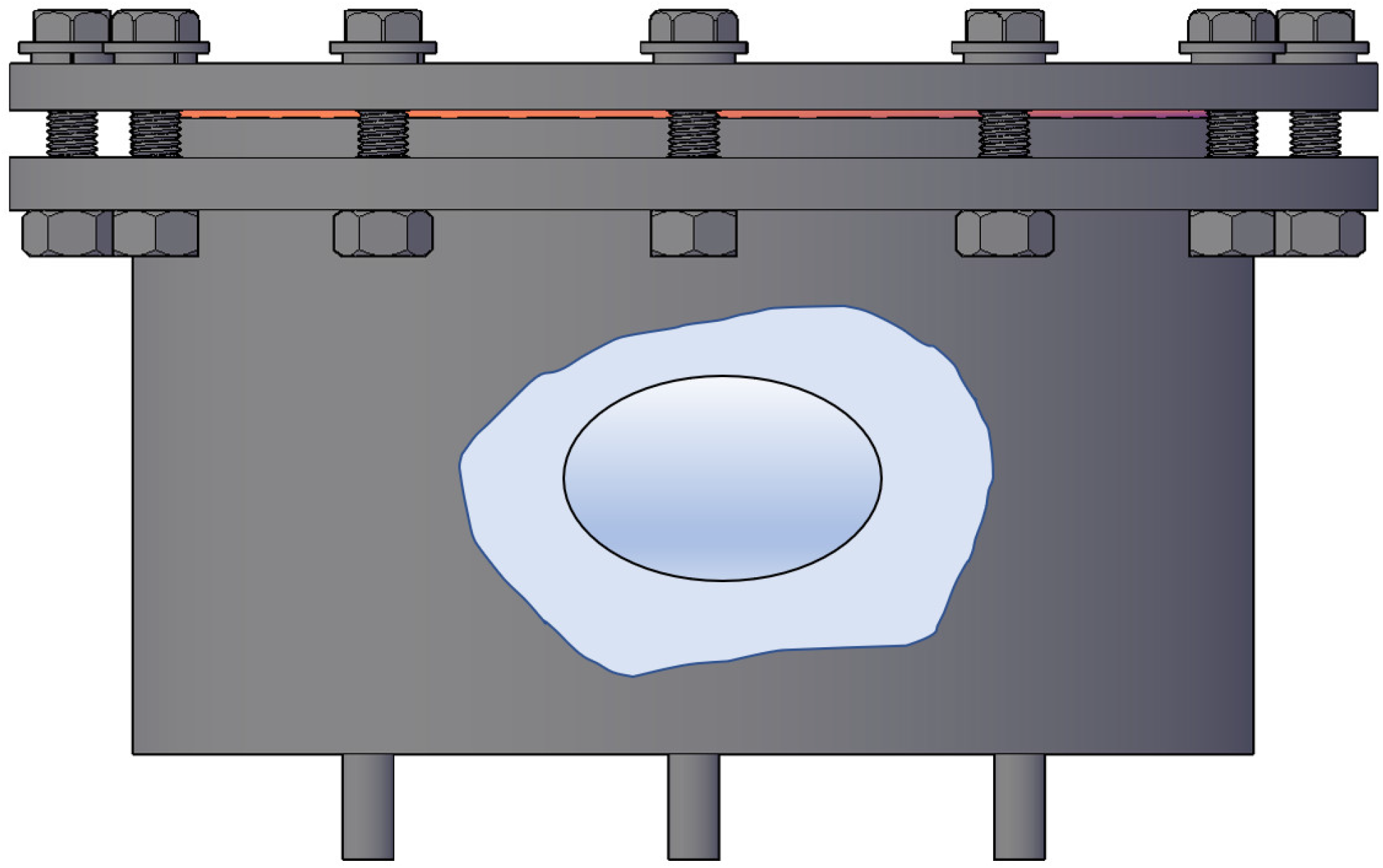
The concept of elimination of ice nucleation on the inner walls of an isochoric chamber

Experiments were performed with a plastic bag with a volume of two liter completely filled with steam distilled water. The plastic LDP bag is placed in the isochoric chamber in such a way that it cannot touch the chamber walls. A 3M saline solution was used to fill the space between the bag and the inner wall of the isochoric chamber following the procedure described in 2.4. Three 48 hours repeats of this experiment have shown that the supercooled fluid did not freeze in this configuration.

### 3.3. *IN VITRO* SUPERCOOLING PRESERVATION OF A PIG LIVER

Experiments with *in vitro* pig livers, were also performed. Three pig livers weighting between 2 and 1.5 kg’s were used from a local butchery. The pig liver is introduced in the 2-liter LDP plastic bag (see Fig 15A). The plastic bag with the liver was filled with physiological saline, carefully to eliminate any air bubbles and sealed. It is placed in a holding plastic bowl inside the container (Fig 15 B). The purpose of the bowl is to separate the plastic container from the walls. A solution of 4 M NaCl was poured in the isochoric chamber with care to not trap any air bubbles. This was done as described in the experimental methods section 2.4 in Materials and Methods. The isochoric chamber was closed and the reactor introduced in the isochoric reactor cooling bath. The temperature was set to – 3 °C and the temperature and pressure monitored in time. (Figure 15 C). Twenty-four, forty-eighth and other 48 hours repeat experiments were performed. There was no ice nucleation in any of the repeats.

**Figure 15.**
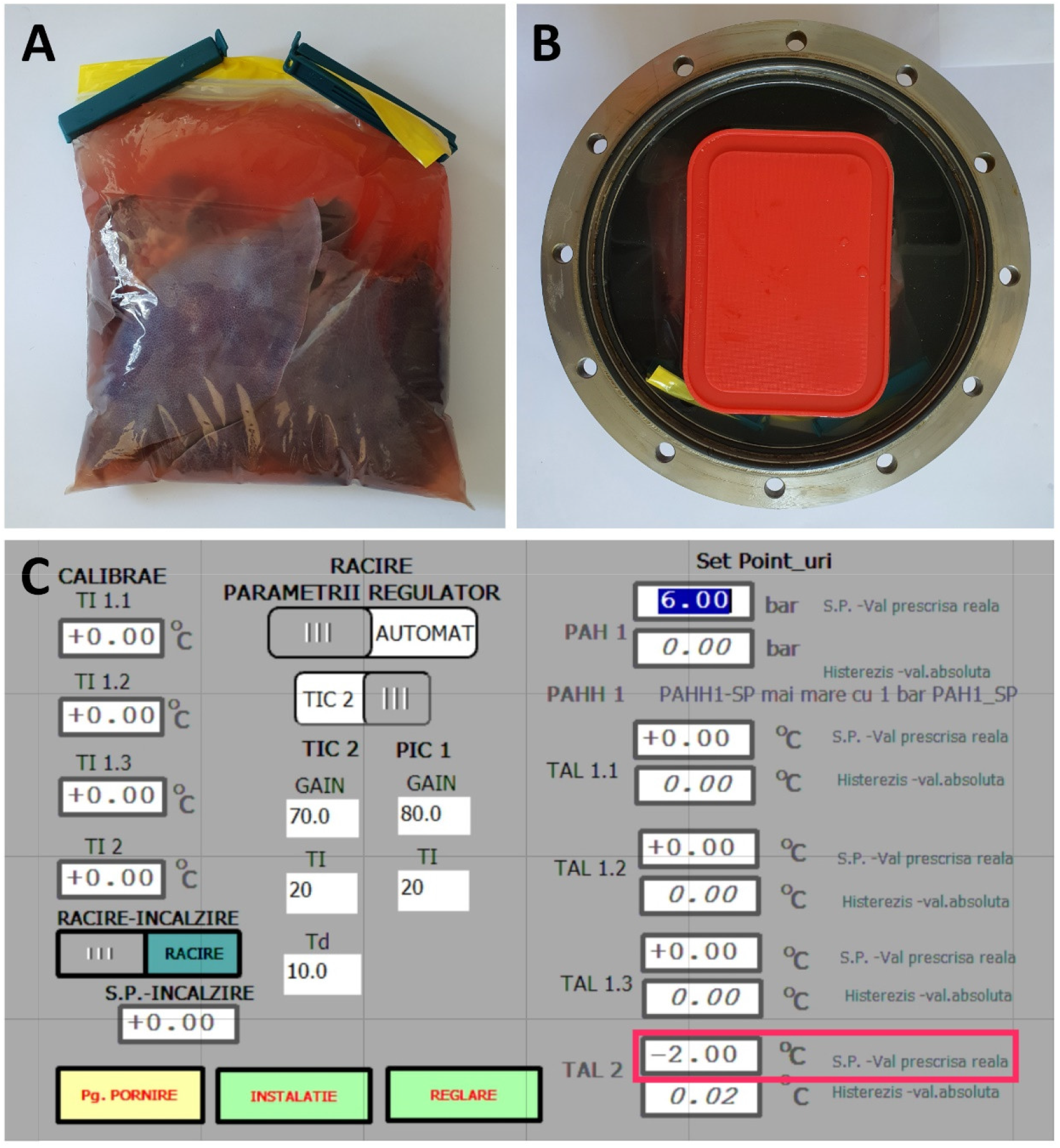
The pig liver inside the 2-liter LDP plastic bag (A); The liver inside the reactor placed between holding plastic boards, to avoid the contact of the bag with the metallic parts of the reactor (B); The temperature set for preservation, in the cooling bath (C)

Figure 16 shows the temperature measured at different locations in the isochoric chamber as a function of time during the isochoric supercooling. No ice nucleation is seen for 48 hours. Figure 17 shows the pressure measured during the supercooling experiment. There is no pressure elevation as that which happens in Figure 11. The experiment was terminated voluntarily.

**Figure 16.**
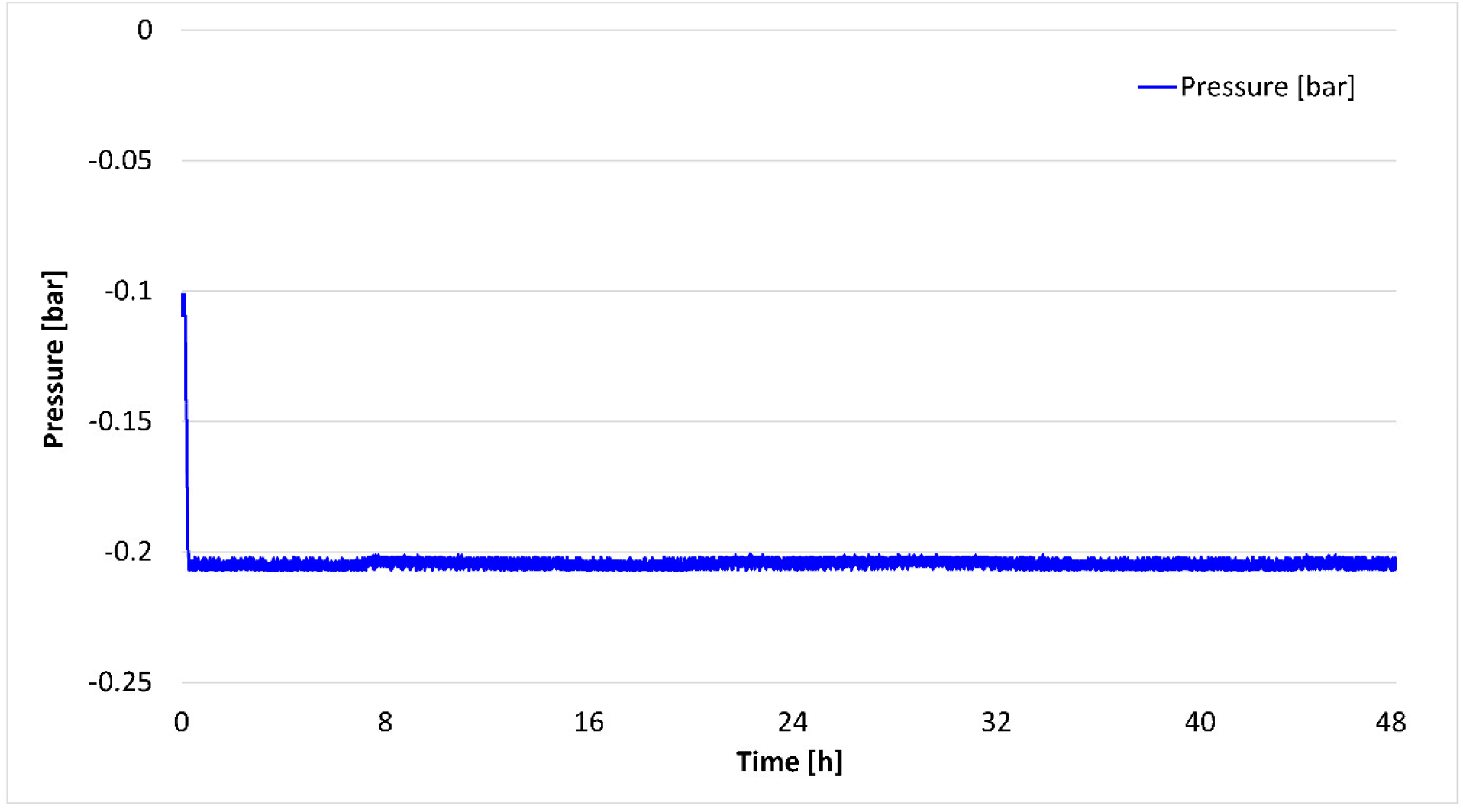
The temperature trace inside the isochoric chamber during the supercooling experiment.

**Figure 17.**
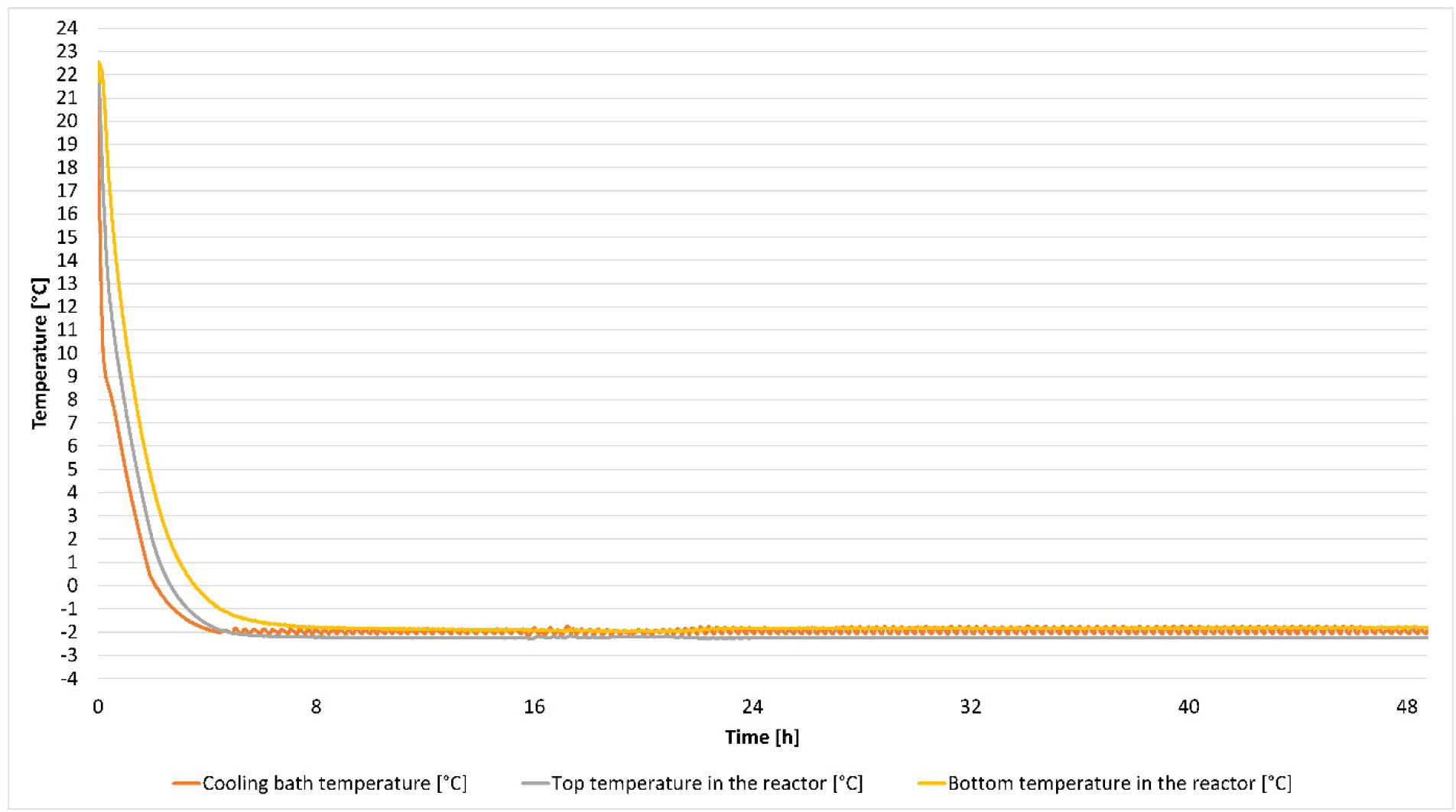
The pressure trace inside the isochoric chamber during the supercooling experiment.

The most important aspect of this paper is to provide researchers in the field of cryobiology with means to perform isochoric experiments with large organs, such as the pig liver. Among other incidentals findings are the observation that pressure can be used to detect ice nucleation. The two-container technique may also aid in inhibition of ice nucleation from the walls of the isochoric chamber.

## 4. SUMMARY

A large 11-liter isochoric chamber for preservation of large organs in an isochoric supercooled state was designed and tested. Details of the design of the chamber are given as well as a demonstration that the chamber is isochoric. Unlike observations with small supercooled isochoric chamber, five repeats show that in this large chamber, ice nucleation of steam distilled water occurs at – 2 °C within less than 12 hours, in all the repeats. An analysis of the results suggests that the ice nucleation begins on the inner walls of the isochoric chamber. A new technique that eliminates the probability of ice nucleation on the walls of the isochoric chamber was developed. The biological matter to be preserved is introduced in a low-density polyethylene bag, filled with the preservation fluid and placed in the center of the isochoric chamber, in such a way that the bag does not touch the walls. The space between the inner walls of the isochoric chamber and the outer walls of the bag were filled with a fluid at such a concentration that it cannot freeze at the storage temperature. The isochoric chamber was filled with this fluid using a technique to eliminate all free air from the isochoric chamber. Experiments with steam distilled water and with *in vitro* pig liver show that the system remained supercooled when the new technique was used. Experiments were terminated at 48 hours of supercooling. This new technology may hold promise for long term preservation of biological organs in a supercooled state, without the use of cryoprotectant additives. It may be suitable for any type of supercooling preservation technology that seeks to avoid nucleation from the chamber walls.

## ACKNOWLEDGEMENT

This work was supported by a grant of the Romanian Ministry of Education and Research, CCCDI - UEFISCDI, project number PN-III-P2-2.1-PED-2019-5409, within PNCDI III.

We also would like to thank Mr. Neacșu Ion, for the continuous effort and support offered to our research group, and we are grateful to SC CRIOMEC SA Galați for the technical aid provided.

## BIBLIOGRAPY

1. Shamseddine I, Pennec F, Biwole P, Fardoun F. Supercooling of phase change materials: A review. Renew Sustain Energy Rev. 2022;158:112172.

2. Luyet B. On the supercooling of water. Phys Rev. 1952;85(4):746–746.

3. Takahashi T, Kakita A, Takashi Y, Yokoyama K, Sakamoto I, Yamashina S. Preservation of rat livers by supercooling under high pressure. Transplant Proc. 2001;33(1–2):916–9.

4. Monzen, K; Hosoda, T; Nagai, R; Monzen, Koshiro; Hosoda, Toru; Hayashi, Doubun; Imai, Yasushi; Okawa, Yasuhiro; Kohro, Takahide; Uozaki, Hiroshi; Nishiyama, Tomoki; Fukayama, Masashi; Nagai R. The use of a supercooling refrigerator improves the preservation of organ grafts. Biochem Biophys Res Commun. 2005;337(2):534–9.

5. Ishine N, Rubinsky B, Lee CY. A histological analysis of liver injury in freezing storage. Cryobiology. 1999;39(3):271–7.

6. Puts, CF; Berendsen, TA; Uygun, K; Bruinsma, BG; Ozer, Sinan; Luitje, Martha; Usta, O Berk; Yarmush M. Polyethylene glycol protects primary hepatocytes during supercooling preservation. Cryobiology. 2015;71(1):125–9.

7. Amir G, Rubinsky B, Horowitz L, Miller L, Leor J, Kassif Y, et al. Prolonged 24-hour subzero preservation of heterotopically transplanted rat hearts using antifreeze proteins derived from arctic fish. Ann Thorac Surg. 2004;77(5):1648–55.

8. Ishine N, Rubinsky B, Lee CY. Transplantation of mammalian livers following freezing: Vascular damage and functional recovery. Cryobiology. 2000;40(1):84–9.

9. Tessier, SN; de Vries, RJ; Toner, M; Tessier Shannon N; de Vries Reinier J; Pendexter Casie A; Cronin Stephanie E J; Ozer, Sinan; Hafiz Ehab OA; Raigani, Siavash; Oliveira-Costa Joao Paulo; Wilks Benjamin T; Lopera Higuita, Manuela; van Gulik, Tho M. Partial freezing of rat livers extends preservation time by 5-fold. Nat Commun. 2022;13(1):DOI: 10.1038/s41467-022-31490-2.

10. Huang H, Yarmush ML, Usta OB. Long-term deep-supercooling of large-volume water and red cell suspensions via surface sealing with immiscible liquids. Nat Commun. 2018;9(1):3201.

11. de Vries RJ, Tessier SN, Banik PD, Nagpal S, Cronin SEJ, Ozer S, et al. Subzero non-frozen preservation of human livers in the supercooled state. Nat Protoc. 2020;15(6):2024–40.

12. Powell-Palm MJ, Rubinsky B, Sun W. Freezing water at constant volume and under confinement. Commun Phys. 2020;3(39):https://doi.org/10.1038/s42005-020-0303-9.

13. Rubinsky B, Perez PA, Carlson ME. The thermodynamic principles of isochoric cryopreservation. Cryobiology. 2005;50(2):121–38.

14. Bilbao-Sainz C, Sinrod AGJ, Dao L, Takeoka G, Williams T, Wood D, et al. Preservation of spinach by isochoric (constant volume) freezing. Int J Food Sci Technol. 2020;55(5):2141– 51.

15. Powell-Palm MJ, Zhang Y, Aruda J, Rubinsky B. Isochoric conditions enable high subfreezing temperature pancreatic islet preservation without osmotic cryoprotective agents. Cryobiology. 2019;86:130–3.

16. Szobota SA, Rubinsky B. Analysis of isochoric subcooling. Cryobiology. 2006;53(1):139–42.

17. Powell-Palm MJ, Koh-Bell A, Rubinsky B. Isochoric conditions enhance stability of metastable supercooled water. ApplPhysLett. 2020;123702:https://doi.org/10.1063/1.5145334.

18. Consiglio AN, Lilley D, Prasher R, Rubinsky B, Powell-Palm MJ. Methods to stabilize aqueous supercooling identified by use of an isochoric nucleation detection (INDe) device. Cryobiology. 2022;106:91–101.

19. Campean S-I, Beschea G-A, Serban A, Powell-Palm MJ, Rubinsky B, Nastase G. Analysis of the relative supercooling enhancement of two emerging supercooling techniques. AIP Adv. 2021;11(5):055125.

20. Powell-Palm, J. M, Charwat V, Charrez B, Siemons B, Healy EK, Rubinsky B. Isochoric supercooled preservation and revival of human cardiac microtissues. Commun Biol. 2021;4:1118(2021).

21. Perez PA, Preciado J, Carlson G, DeLonzor R, Rubinsky B. The effect of undissolved air on isochoric freezing. Cryobiology. 2016;72(3):225–31.

22. Preciado JA, Rubinsky B. Isochoric preservation: a novel characterization method. Cryobiology [Internet]. 2010 Feb [cited 2012 Apr 18];60(1):23–9. Available from: http://www.ncbi.nlm.nih.gov/pubmed/19559692

